# Effective NGN2-based neuronal programming of hiPSCs in an automated microfluidic platform

**DOI:** 10.1101/2022.10.13.512042

**Authors:** S Angiolillo, S Micheli, O Gagliano

**Affiliations:** Department of Industrial Engineering (DII), University of Padova, Padova, Italy; Venetian Institute of Molecular Medicine (VIMM), Padova, Italy

**Keywords:** Lab-on-a-chip, Neuronal micro-tissue, NGN2-programming, Tissue engineering, High-throughput screening

## Abstract

Neurodegenerative diseases represent an increasing health burden, with a worrying lack of models recapitulating the hallmarks of the pathology.

Recently, lab-on-a-chip technology has opened new reliable alternatives to conventional *in vitro* models able to replicate key aspects of human physiology. For instance, microfluidics allows to mimic the extracellular accumulation of misfolded proteins in the finely controlled microenvironment, thanks to the intrinsic high surface-area-to-volume ratio.

Automated microfluidic platforms offer advantages in implementing high-throughput, standardized and parallelized assays, suitable for drug screenings and developing new therapeutic approaches in a cost-effective way.

However, the major challenges in the broad application of automated lab-on-a-chip in biological research are the lack of production robustness and ease of use of the devices.

Here, we present an automated microfluidic platform able to host the rapid conversion of human induced pluripotent stem cells (hiPSCs) into neurons via NGN2 viral programming in a user-friendly manner.

The design of the platform, built with multilayer soft-lithography techniques, shows easiness in the fabrication and assembly thanks to the simple geometry and experimental reproducibility at the same time.

The all operations are automatically managed from the cell seeding, medium change, doxycycline-mediated neuronal induction, and selection of the genetically engineered cells, to the analysis, including immunofluorescence assay.

Our results show a high-throughput, efficient and homogenous conversion of hiPSCs in neurons in 10 days showing the expression of mature marker MAP2, and calcium signaling. The neurons-on-chip model here described represents a fully automated loop system able to address the challenges in the field of neurodegenerative diseases and improve current preclinical models.

## Introduction

Neurodegenerative diseases, which involve the progressive loss of neuronal cell function, represent a major global health challenge impacting patients as well as public healthcare system^1,2^.

Although extensive research efforts aim to develop new disease-modifying therapeutics, there is no significant progress resulting from the high failure rates of developed models^3–5^. The biggest limitation in neurodegenerative disease research is the lack of *in vitro* human models recapitulating the pathological hallmarks^6^: for instance, in conventional neuronal cultures the level of misfolded proteins is insufficient to form aggregates, and the extracellular ones are also washed out by medium change. Moreover, current protocols for *in vitro* generation of human neurons are still often lengthy, expensive, complex and yield heterogenous neural populations^3^.

Recently, lab-on-a-chip technology has opened new reliable alternatives to conventional *in vitro* models able to replicate key aspects of human physiology. These microfluidic devices have complete control over the cellular environment with high spatial and temporal resolution and precisely defined geometries^7,8–11^.

The intrinsic property of high surface-to-volume ratio provides fast and cyclic stimuli^12^,a timely controlled supply of nutrients, accumulation factors^13,14^, and dynamic analysis of output in response to specific cues^12,15^.

The accumulation of endogenous secreted factors in the confined microenvironment^16^ is the key point to choose microfluidic *in vitro* models to mimic neurodegenerative disorders^17^, as they are characterized by appearance of intracellular and extracellular deposition of toxic proteins in the brain, which progressively lead to the neuron death^11^.

Neurons-on-chip *in vitro* models mainly rely on the use of primary cell lines^18–22^ and embryonic stem cells (ESCs)-derived neurons^23,24^. They are able to form complex neuronal networks within the microfluidic environment, where to study neurite growth upon exposure to biochemical gradients, cell-cell synaptic interactions, and the formation of the blood-brain-barrier. However, primary cell lines derived from rodent embryos are terminally differentiated, mature neurons, which limits their application to a single assay, and can show species-specific differences in cell behavior. The use of genetically modified human embryonic stem cells (hESCs) allowed to obtain an unlimited source of human cells to differentiate into neurons, but their isolation is limited by ethical concerns and cannot be derived from adult patients^25^.

Advances in stem cell technology, such as human induced pluripotent stem cells (hiPSCs)^26^, offer a great potential to develop patient-specific preclinical models: hiPSCs-derived neurons can model both the most common sporadic forms and the least common familial forms of neurodegenerative diseases, without any genetic modification, and thus they represent promising tools for personalized drug screenings and for the discovery of new therapeutic targets.

To this aim, it is fundamental to rely on a standardized protocol which combines high efficiency of neuronal conversion and short generation time.

Neurogenin2 (NGN2) is a pro-neural transcription factor able to obtain a homogeneous population of cortical neurons from pluripotent stem cells in a time-effective way^27^.

This differentiation strategy in combination with microfluidics was reported only in few papers^28,29,30^, showing the possibility to obtain human cortical neurons, expressing mature pan-neuronal marker MAP2, in a controllable microenvironment in just one week.

The limitations to apply such models to preclinical studies are the low potential of parallelization, throughput and standardization, key elements required from pharmaceutical pipelines^8^.

Automated microfluidic platforms offer advantages in implementing high-throughput, standardized and parallelized assays, suitable for drug screenings and developing new therapeutic approaches in a cost-effective way^31^.

The major challenges in the broad application of automated lab-on-a-chip in biological research are the lack of production robustness and ease of use of the devices^9^.

In this work, we presented an automated microfluidic platform composed by 8 chambers which ensure parallelization of the experiments, high throughput of tested conditions and the experimental reproducibility. The design of the platform, built with multilayer soft-lithography techniques, shows easiness in the fabrication and assembly thanks to the simple geometry and experimental reproducibility at the same time.

To our knowledge there are no reports showing neuronal programming of hiPSCs using NGN2 in an automated microfluidic platform.

Our results show a high-throughput, efficient and homogenous conversion of hiPSCs in neurons in 10 days showing the expression of mature marker MAP2, and calcium signaling. The neurons-on-chip model here described represents a fully automated loop system useful to address the challenges in the field of neurodegenerative diseases and improve current preclinical models by decreasing the time and costs and allowing dynamic studies.

## Materials and Methods

### 2.1 Description of microfluidic chip design

The automated microfluidic chip was produced by standard photo- and soft-lithography techniques^32^.

It is composed of two polydimethylsiloxane (PDMS, Sylgard 184, Dow Corning) layers: a flow layer, for hosting the cellular model, and a control layer (Fig. 1A).

**Fig. 1.**
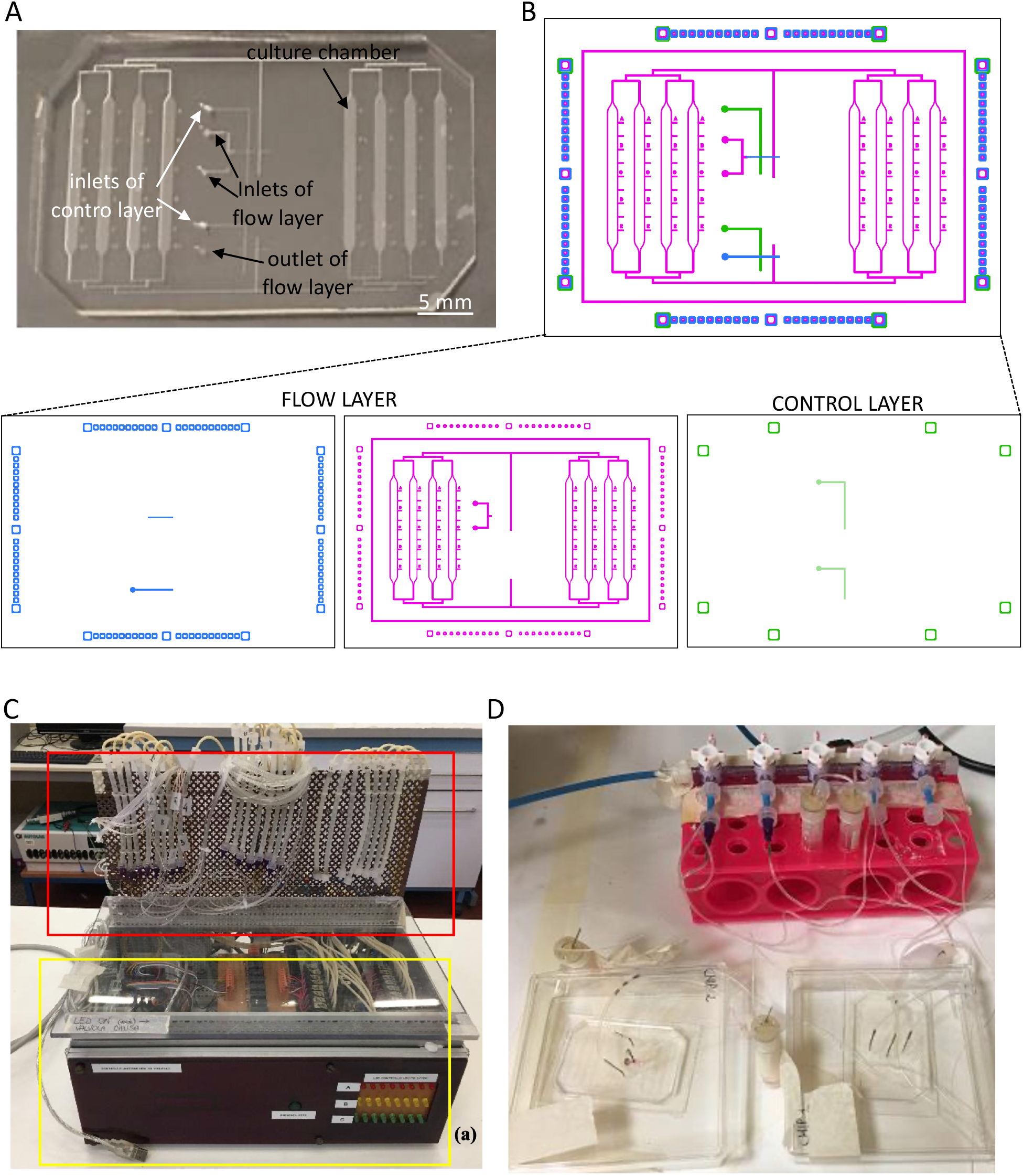
Experimental setup of the multilayer microfluidic platrform. A) Microfluidic device developed after the process of Replica molding. B) Multilayer photomask of the microfluidic device disegned using Autocad. In the detail below, the flow layer consists in two layers, one in violet made with negative photoresist and another in blue made with positive photoresist; the control layer, in green, is made with negative photoresist. C) Hardware instrumentations used to automate and control the multilayer platform. D) Equipment for fluid distribution. Pressurized vials are connected to the inlets of the platform by Teflon tubes.

The design of the two photomasks was performed using the software AutoCAD® (Fig. 1B). The device includes 8 independent and parallel culture chambers, connected to the two inlets and one outlet by a network of flow channels (1.7 mm width x 0.25 mm height); each culture chamber (25.8 mm length x 1.7 mm width x 0.25 mm height) presents a surface of 40 mm^2^, with a volume of 11 μL. The control channels have a width of 200 μm and a height of 35 μm.

### 2.2 Control layer design

A silicon wafer of 4 inches was cleaned with acetone (Sigma-Aldrich), methanol (Sigma-Aldrich) and distilled water in order to promote the photoresist adhesion on the wafer in following steps. The wafer was dried using compressed air and then is placed on a hot plate at 105°C for 20 minutes to remove humidity. The wafer was inserted in the spin coater (WS-650-23NPP, Laurell) to deposit one layer of SU-8 2050 negative photoresist (Microchem) to obtain a thickness of 35 μm. After a Soft Bake heating at 95°C for 6 minutes, the wafer coated was patterned by UV exposure at an exposure dose of 160 mJ/cm^2^ using the designed photomask (Fig. 1B).

After a post-exposure bake at 95°C for 6 minutes, the wafer was developed using propylene glycol monomethyl ether acetate (Sigma-Aldrich) for 5 minutes, then rinsed with isopropyl alcohol (Sigma-Aldrich) and dried with compressed air. Hard Bake step was done putting the wafer on a hot plate at 65°C for 10 minutes and then setting a ramp of 120°C/h for 2 hours to reach the final temperature of 160°C.

### 2.3 Flow layer design

A silicon wafer of 4 inches was cleaned and dried following the same procedure described for the control layer. The flow layer needs to be done by two different photoresists: the positive resist, was used to build round shape channels ensuring a proper closing of the valve, whereas the negative resist was used for the other components of the flow geometry. In our work we build the microvalves according to the push-up geometry.

For the first layer of positive resist, the wafer was inserted in the spin coater (WS-650-23NPP, Laurell) to deposit one layer of SPR-220-7 positive photoresist (Rohm and Haas, Dow Corning) was distributed and spin-coated over the wafer in order to obtain a thickness of 15 μm. After a Soft Bake heating at 90°C for 10 minutes, a second layer of the same photoresist was spun over the first one to obtain a finale thickness of 50 μm. A Soft Bake was done at 90°C for 60 minutes. Then, the wafer was maintained in the dark for 3 hours to rehydrate the photoresist. The wafer coated was patterned by UV exposure at an exposure dose of 200 mJ/cm^2^ using the positive photomask of the flow layer (Fig. 1B).

After a post-exposure bake at 95°C for 15 minutes, the wafer was developed in a solvent, composed by water (97-98%) and tetramethylammonium hydroxide (2,45%), (Microposit MF-319, Microchem) to dissolve not cross-linked polymer for 15 minutes, it was rinsed with distilled water to stop the development and dried with compressed air. As last step the wafer was cooked on a hot plate at 65°C for 10 minutes and then of 10 °C/h for 15 hours to reach the maximum temperature of 190°C.

A second layer of SU-8 2100 negative photoresist (Microchem) was spin-coated over the wafer to obtain a thickness of 250 μm. After a Soft Bake heating at 95°C for 45 minutes, the wafer was patterned by UV exposure at an exposure dose of 350 mJ/cm^2^ using the negative photomask of the flow layer (Fig. 1B). After a post-exposure bake at 95°C for 15 minutes, the wafer was developed using propylene glycol monomethyl ether acetate (Sigma-Aldrich) for 17 minutes.

At the end the wafer was rinsed with isopropyl alcohol (Sigma-Aldrich) and then dried with compressed air. Hard Bake step was done putting the wafer on a hot plate at 65°C for 10 minutes and then setting a ramp of 120°C/h for 2 hours to reach the final temperature of 160°C.

### 2.4 Device assembly

The two layers of the chip were made through replica molding in PDMS (Sylgard®184, Dow Corning), by mixing a curing agent (cross-linker) and a base (siloxane) in a weight ratio of 1:10. For the flow layer, a 4 mm of PDMS layer was poured on the silicon mold, and once baked at 60°C for 2 hours, it was peeled off and punched in correspondence to the two inlets and the outlet with a dispense tip (Nordson EFD) having external diameter equal to 0.91 mm. Whereas the control layer was made by spin-coating a layer of PDMS over the wafer to obtain a thickness of about 80 μm, that was baked with a heating ramp of 40°C/h for 2 hours to reach 80°C.

For building the chip, the flow layer was coupled with plasma treatment over the control layer, without peeling it from the mold. The alignment marks simplified the procedure of alignment of the two layers of the device.

Once the two layers were bond each other, the multilayer PDMS block was cut, peeled off, punched to create the inlets for the control layer and attached over the glass slide with plasma treatment (Fig. 1A). The device was sterilized by autoclaving before use.

### 2.5 Automation set-up

Microfluidic chip was equipped with an integrated system of remotely controlled pneumatic valves that distributed medium with precise timing into the 8 culture chambers.

In Fig. 1C is reported the hardware instrumentations used to automate and control the multilayer platform.

The main system is mounted inside an aluminium box in order to make easier the moving of the system in the laboratory. The main system is composed by three parts: i) Electronic unit for the generation of the signal (Fig. 1C, in yellow); this part is connected to an external computer to generate the digital input. ii) Solenoid valves (Fig. 1C, in red): each of them is connected to a tube (PTFE, Cole Palmer) that will be filled with distilled water and connected to the control inlets of the microfluidic platforms; each tube ends with a needle to be easily inserted into the holes. iii) Pressure regulator: the pressure of solenoid valves can be easily controlled and changed by a pressure regulator (CDK Corporation).

All the on-chip valves are driven by solenoid valves (24V, CDK Corporation) which are controlled by an electronic unit for the generation of digital signal (NI-9472, National Instruments) connected to the USB port of a computer.

LabVIEW® is needed to create an interface for controlling the instrumentation from a computer. Each solenoid valve can switch on-chip valve from atmospheric pressure (on-chip valve open) to the pressure that permit the closure of the on-chip valve through a pressurized liquid.

Last, the fluid distribution equipment (Fig. 1D) is composed by pressurized vials, connected both to the pressurized air line and to the inlets of microfluidic platform.

### 2.5 Cell culture integration

During experiments the microfluidic chip was placed in a dish, surrounded by a water bath to reduce medium evaporation.

After connecting the control tubes and flow tubes and performing a debubbling step to avoid bubble formation, we coated the glass surface to promote the cell adhesion with 25 μg/ml of Fibronectin (Thermo Fisher Scientific) for human fibroblasts BJ culture, and 2.5% (vol/vol in DMEM) Matrigel Reduced Factor (MRF) for hiPSCs. The coating solution was inserted into a pressurized vial and injected through the inlet of the platform and incubated for 1 hour at room temperature.

Finally, the cell suspension with the desired density, 20-40-100 cells/mm^2^ for BJ fibroblasts and 125-250-500 cells/mm^2^ for hiPSCs infected with TetO-Ngn2-T2A-Puro lentiviral vector (hiPSCs-NGN2), was placed inside a pressurized vial and injected through the inlet in the microfluidic platform.

The BJs are seeded with DMEM high glucose supplemented with 10% FBS, whereas the hiPSCs with TeSR-E8 medium (Stem Cell Technologies) supplemented with 10 μM of ROCK inhibitor (Y-27632, Sigma-Aldrich)

After 12 hours from the cells seeding, the culture medium was automatically changed every 12 hours.

### 2.6 Neuronal differentiation on chip

For the neuronal differentiation experiment, we seeded the hiPSCs-NGN2 (Day −1) with the highest density among the three tested (500 cell/mm^2^), after assessing that it was the most efficient one (data not shown).

The day after seeding of hiPSCs-NGN2 (Day 0), the culture medium is replaced with TeSR-E8 medium supplemented with Doxycycline (2 ng/ml, Sigma-Aldrich) to induce TetO gene expression and Puromycin Dihydrochloride (1 ng/ml, Thermo Fisher Scientific) to select only the genetically engineered cells.

On Day 1, medium was switched to N2B27 medium and changed every 12 h per day untill Day 4. From Day 5 untill Day 10, medium was replaced every 12 h with iN medium, consisting in N2B27 medium supplemented 10 ng/ml of Brain-derived neurotrophic factor (BDNF) and 10 ng/ml of Neurotrophin-3 (NT3).

### 2.6 Immunofluorescence assay

Cells were fixed using 4% paraformaldehyde solution incubated for 10 minutes at room temperature. Blocking solution (PBS supplemented with 0.1% triton-X-100 (Sigma-Aldrich, #93426) and 5% horse serum (Sigma-Aldrich)) was then added to each channel and incubated for 1 hour at room temperature. After 1 hour all the chambers were automatically washed for three times with a solution of PBS-/-with 0.1% of Triton X-100 (PBST).

Cells were incubated with primary antibodies diluted in blocking solution overnight at 4°C. Cells were then washed three times with PBST, incubated for 1 hour at room temperature with secondary antibodies diluted in blocking solution, washed again three times in PBST and mounted with mounting solution. The following antibodies were used for our analysis: rabbit anti-NGN2 (D2R3D) (1:100, 13144S Cell Signaling), mouse anti-OCT4-3/4 (C-10) (1:200, sc-5279 Santa Cruz Biotechnology) and mouse anti-TUBULIN β 3 (βIIITUBB) (1:5000, 801202 BioLegend) and MAP2 (1:10000, Ab5392 Abcam).

Appropriate Alexa Fluor 488- and Alexa Fluor 594-conjugated secondary antibodies (1:200, Jackson ImmunoResearch) were used. Nuclei were counterstained with Hoechst (1:5000, H3570 Life Technologies).

Cell viability assay (Live/Dead kit, Thermo Fisher) was performed on microfluidic differentiated cells. Cells were washed with DMEM high glucose, incubated with 3 μM ethidium homodimer-1 (stains dead cells in red), 3 μM calcein AM (stains live cells in green) and 4 μM Hoechst (stains cell nuclei in blue) for 45 minutes at 37°C, washed with DMEM and analyzed at fluorescence microscope.

Pictures were taken with Leica DMI6000 B fluorescence microscope.

### 2.7 Fura-2 Ca^*2+*^ imaging

[Ca^2+^]_c_ was measured by ratiometric analysis using acetoxy-methyl-ester Fura-2 (Fura-2/AM; F1221 Thermo Fisher) according to the manufacturer’s instructions. Cells at Day 10 in the microfluidic device were incubated with 1 μM Fura-2 in iN medium for 30 minutes and washed three times with fresh iN medium. Image processing was performed using an inverted microscope (Olympus IX81) with 40X UV objective lens. For Fura-2 excitation, sequential illuminations at 340 nm and 380 nm were delivered every 3 seconds. Emission of Fura-2 was detected at a wavelength of 510 nm. Ratios of emission fluorescence following excitation at 340 nm and 380 nm were recorded and calculated by AIW. For every recording, 3-5 cells were randomly chosen, and soma were analysed. After 6 minutes from the beginning of the image acquisition, iN medium supplemented with 65 mM KCl was administered to measure cell depolarization.

## Results and Discussion

### 3.1 Characterization of fluid dynamics in the system

Before integrating the cellular model, the designed microfluidic platform was characterized for the fluid-dynamic behaviour.

After assembling and connecting all the tubes to pressure lines, the two inlets of the device was connected to the pressurized vials containing two different dyes, one yellow and one red, A video was recorded while the dyes were flowing into the chambers, during a sequential flow change from one dye to another, as reported in Fig. 2A. This experiment allowed to verify the presence of pressure drop in the microfluidic circuit that could affect the media change of the culture chambers.

**Fig. 2.**
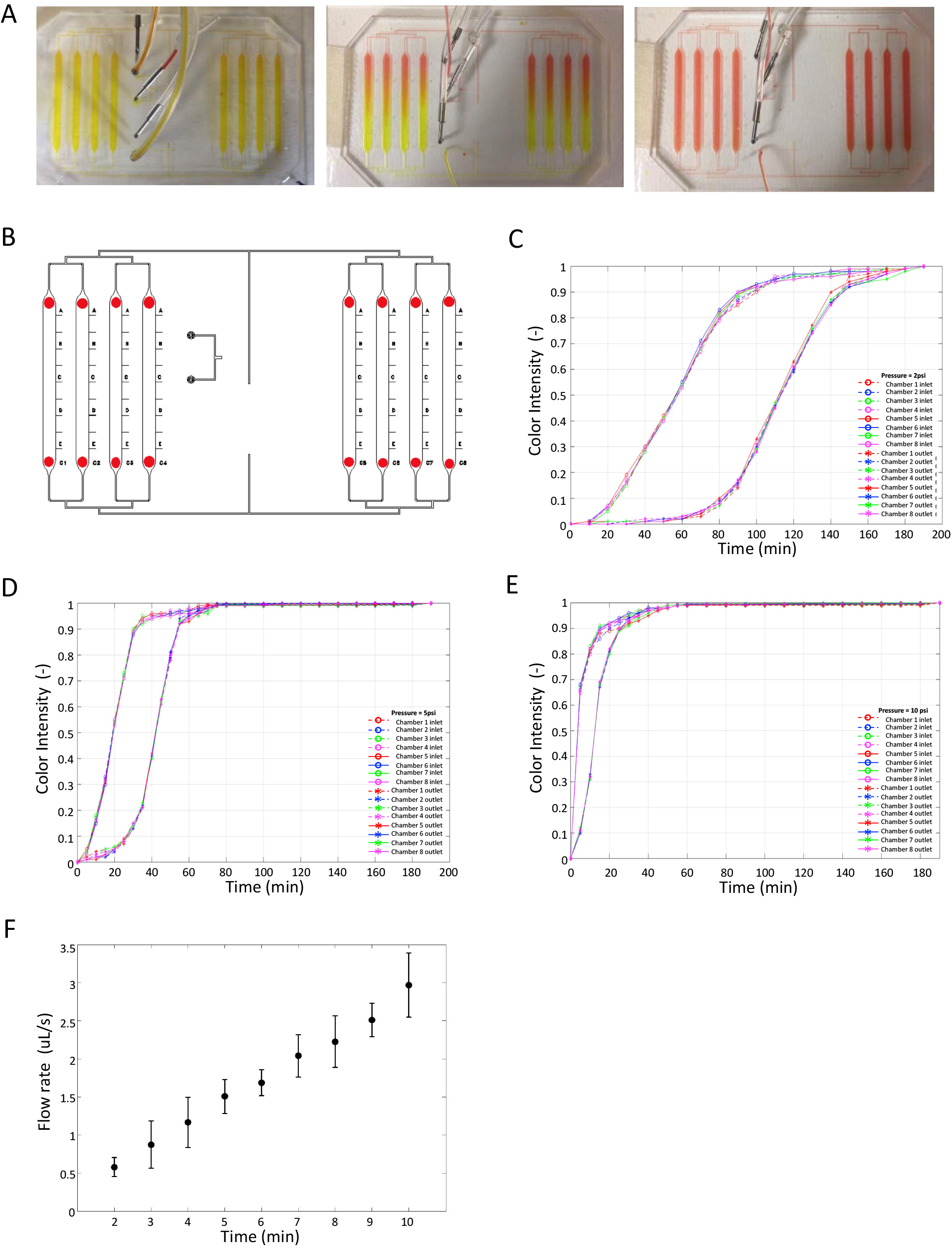
Fluid-dynamics behavior of the multilayer platform. A) Time lapse of medium change images of a representative experiment. Left, initial situation where all the chambers are filled with yellow dye; middle, intermediate situation where the chambers are filled with dye colorant; right, final situation where all the chambers are filled with red dye. B) Placement of the 16 ROIs placed near the inlet and the outlet of each chambers. The ROIs are the small red circle. C),D), E) Colour intensity [-] vs time [s] of the ROIs placed near the inlet and the outlet of all the chambers of the microfluidic platform for a flow pressure of 2, 5 and 10 psi respectively. F) Volumetric flow rate [μl/s] vs pressure [psi] in the microfluidic device.

Two ROIs were placed in correspondence to the inlet and outlet of each chamber, as reported in Fig. 2B, to investigate the presence of diffusion phenomena and flow delay during the medium change due to the imperfections in opening/closing of the micro-valves.

Assuming yellow as value 0 and red as value 1, we characterized the fluid dynamics values by quantifying the colour intensity of the ROIs defined in three different conditions of flow pressure, 2, 5 and 10 psi, respectively (Fig. 1C-E).

In all conditions, profiles started from a value 0, corresponding to the yellow dye, and increased toward the value 1 that was completely reached at medium change accomplished with red dye, with a trend slope and the time of medium change depending on the flow pressure as expected.

From the video we also determined the volumetric flow rate at the different flow pressures imposed from 2 to 10 psi, by measuring the time that dye took to reach a specific length in the channel, and then multiplied by the cross-section area of each channel^33^.

As Fig. 2F shows, the minimum flow rate is 0.58 μL/s ± 0.12 achieved with a pressure of 2 psi whereas the maximum value is 2.97 μL/s ± 0.42, corresponding to a pressure of 10 psi. Moreover, a linear relationship between the volumetric flow rate and the height was observed, in accordance with the Hagen–Poiseuille’s equation.

Last, to control the mechanical resistance of platform and of microvalves for long period of time (30 days), we performed the leakage test at 2, 5 and 10 psi.

According to these results, we selected 5 psi as the working flow pressure because guaranteed a fast medium change (80 seconds, Fig. 2D) with a flow rate of 1.5 μL/s ± 0.23, without any lamination phenomenon of the platform.

### 3.2 Integration of cellular model on chip

The biological validation of the microfluidic platform has been done first using human foreskin BJ fibroblasts. We automatically seeded three different density of cells, 20 cells/mm^2^, 40 cells/mm^2^ and 100 cells/mm^2^ (Fig. 3A), comparing the growth rate to conventional 2D culture (data not shown). The seeded cells were homogenously distributed, and they proliferated without significant differences along the chamber.

**Fig. 3.**
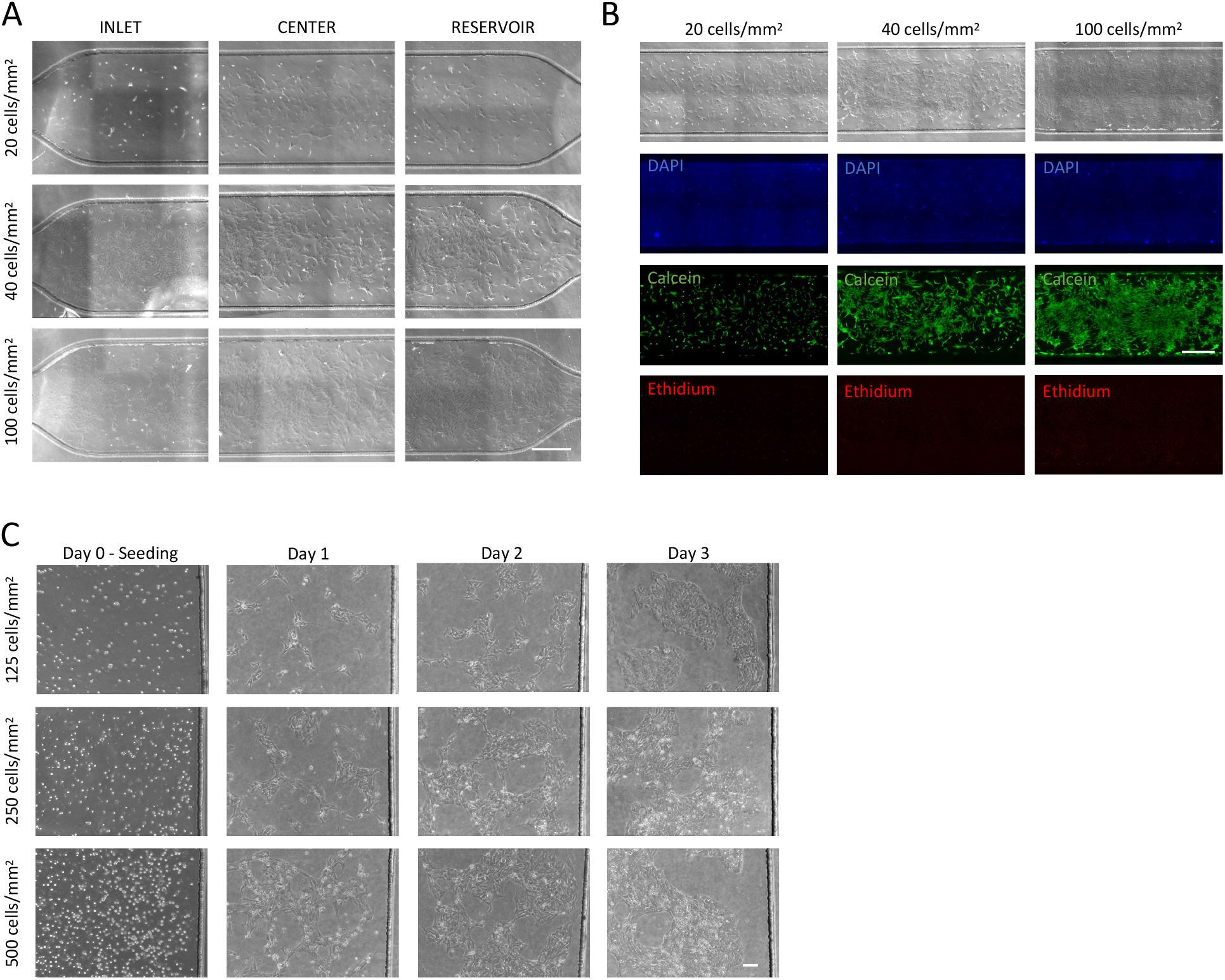
Biological validation of the multilayer platform. A) Representative inlet/center/reservoir regions of the culture chambers after 20 days of fibroblasts culture. The initial seeding densities of fibroblasts were: (a) 20 cells/mm^2^; (b) 40 cells/mm^2^; (c) 100 cells/mm^2^. Scale bar 50 um. B) Cell viability (Live/Dead assay) inside the chambers at day 20 for all the 3 different cells densities: (a) 20 cells/mm^2^; (b) 40 cells/mm^2^; (c) 100 cells/mm^2^. Scale bar 50 um. C) Rappresentative pictures of iPSCs seeding and growth inside the culture chambers with three different seeding densities: (a) 125 cells/mm^2^; (b) 250 cells/mm^2^; (c) 500 cells/mm^2^. Scale bar 100 um.

After 20 days of culture, the viability of the cells was assessed using a commercial live/dead assay kit (Fig. 3B), according to the recommended protocol. The three densities tested reached different cell confluence, depending on the initial density, and showed long-term viability on the multilayer platform.

Since these results encouraged the use of the platform developed with a cellular model, we tried to integrate a hiPSC culture. In this case, due to the smaller dimension of stem cells comparing to mesenchymal ones, we increased the initial seeding density, performing a comparison among 125 cells/mm^2^, 250 cells/mm^2^ and 500 cells/mm^2^.

All cell densities were automatically integrated into the chambers through the microfluidic circuit and were able to proliferate in the confined environment.

In this case the culture on chip was monitored only for three days (because of higher proliferation they reach the confluence early), given that the purpose of the platform was to develop an efficient model of neurons on chip obtained from hiPSCs in a time-saving way.

### 3.3 Neurons on chip

To develop a neuron model on chip, we used a hiPSC line previously infected with TetO-Ngn2-T2A-Puro lentiviral vector, that allows an efficient neural conversion of stem cells by exposure to doxycycline^27^.

The differentiation scheme (Fig. 4A, top) consisted in: seeding (Day −1), doxycyclin addition (Day 0), neuronal induction with doxycyclin and puromycin selection (Day 1 - Day 4), and neural maturation (Day 5 - Day 9).

**Fig. 4.**
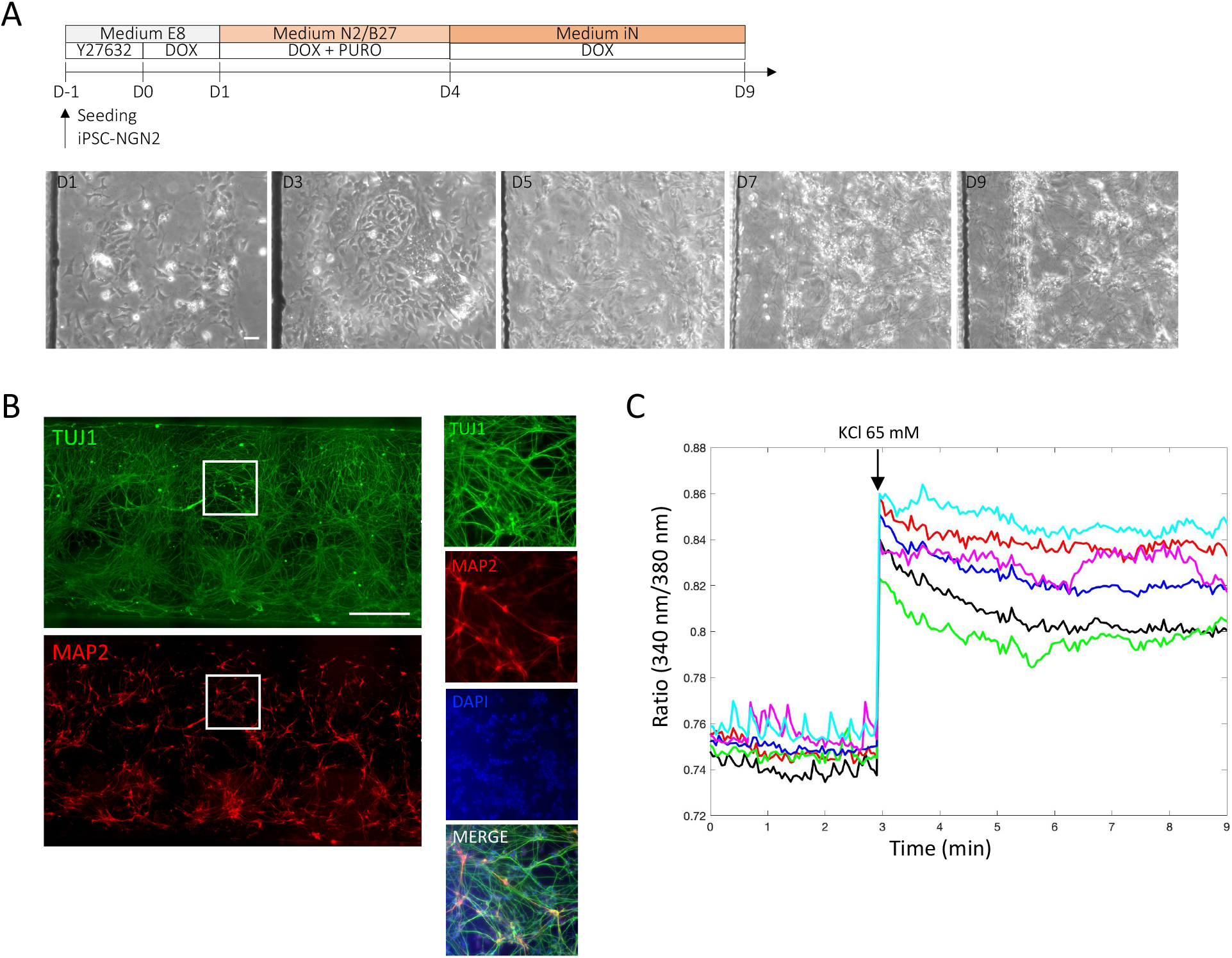
Neuron on chip model. A) Top, schematic representation of neuronal induction via Doxycyclin-mediated NGN2 overexpression and Puromycin selection; bottom, representative pictures showing the morphological evolution from iPSC to neurons inside the culture chamber. Scale bar: 100 μm. B) Representative immunofluorescence images showing neuronal markers TUJ1 (green) and MAP2 (red). Scale bar 50 um. C) Graph showing voltage-gated Ca^2+^ responses in six iPSC-NGN2-derived neurons at day 9 by exposing to 65 mM KCl. Neurons were loaded with Fura-2.

After one day from doxycyclin administration, the cells started their phenotype change to a more elongated morphology and gradually assumed a neuronal appearance, as reported in Fig. 3A. From Day 7, the cells showed the typical morphology of neurons: a central axon and several processes extending towards and away from the cell body.

After 10 days of NGN2 overexpression we observed an efficient hiPSC differentiation toward neural fate with high purity. The neural on chip model obtained showed neuron-specific phenotype expressing TUJ1 and mature neuronal markers (MAP2) along the channel (Fig. 4B).

The model on chip was also characterized the intracellular calcium in real time. As shown in Fig. 4C, after loading the neurons with Fura-2, we depolarized with 65 mM KCl to activate voltage gated calcium channels. As expected from a responsive cellular model, the stimulus applied caused an initial increase in the Fura-2 ratio compared to the steady state, followed by a prolonged and sustained plateau.

## Discussion

In this work we developed an automated multilayer microfluidic platform able to guide the neuronal differentiation of hiPSCs via NGN2 over expression.

The platform developed satisfied the major challenges that slow down the automated lab-on-a-chip model spread in biological research, due to the fabrication complexity requiring a high level of expertise.

The automated lab-on-chip platform displays a simple design, ensuring high throughput at the same time. The 8 parallel culture chambers, with a 11 uL volume, allow the possibility to perform high number of replicates in parallel with a consistent reduction of the cost, in a completely operator-independent manner.

The low number of microvalves and the presence of alignment marks simplified the procedure of fabrication and coupling of the different layers, reducing the rejection rates of the assembled chips^34^. Moreover, the presence of only 2 valves dramatically reduced lamination episodes and guarantees productions of durable and resistant platforms for long term experiments.

From the biological point of view, the device displayed a good biocompatibility with three different cellular models, human fibroblasts, hiPSC and hiPSC-derived neurons (Fig. 3-4). Indeed, the long-term availability on chip represents a great advantage to use the platform for long term studies. We did not observe significant differences in cell density along the channel, both from the seeding to the day-by-day maintenance, thanks to the automated flow distribution allowing a precise control of the microenvironment.

With this positive evidence, we developed a fully automated neurons model on chip via NGN2-overexpression, efficient both in time and in phenotype conversion, never shown until now.

10 days of NGN2 induction were sufficient to instruct hiPSCs toward a neural fate; human neurons, which occupied all the microfluidic chamber section, showed a markable expression of neuron-specific phenotype TUJ1 and mature neuronal markers MAP2 (Fig. 4B) and activation of voltage gated calcium channels (Fig. 4C).

Although our results in terms of neural differentiation efficiency are comparable with other works already published^28,29,30^, here we designed an automated platform able to overcome the limitations of the manual-handling and low parallelization^8^.

Our platform offers the possibility to perform high-throughput studies (8 replicates per condition) remotely controlled, with high precise control of the culture microenvironment and maximizing the robustness of the results.

The low level of automation complexity allows to put several platforms in parallel (Fig. 1D), providing the possibility to perform different experimental conditions, one per chip, with a robust number of technical replicates.

Moreover, the strategy used to obtain human neurons, based on NGN2-overexpression in hiPSCs, allowed to rapidly and cost-effectively produce high-yield patient-specific neurons on chip.

## Conclusion

The NGN2-based neuronal programming of hiPSCs in automated microfluidic platform presented in this work,

Our work represents the first attempt of NGN2-based neuronal programming of hiPSCs in automated microfluidic platform. Thanks to the highly reproducible environment controlled by an automated system we were able to develop a standardized, parallelized, miniaturized and patient-specific neuronal model.

This platform matches the needs of pharmaceutical companies R&D of improving speed, cost and reproducibility for large-scale preclinical studies^35,36^.

This platform can provide dynamical screens with drug cocktails and signaling molecules, by precisely tuning concentration, timing, and duration of delivery in an automated fashion, with minimal training and amounts of reagents.

## Acknowledgements

This project is supported by the University pf Padova under the 2019STARS Grants programme. Dr. Cecilia Laterza (University of Padova) kindly provided NGN2 lentivirus.

## References

1. Nichols, E. et al. Estimation of the global prevalence of dementia in 2019 and forecasted prevalence in 2050: an analysis for the Global Burden of Disease Study 2019. Lancet Public Heal. 7, e105–e125 (2022).

2. Strianese, O. et al. Precision and personalized medicine: How genomic approach improves the management of cardiovascular and neurodegenerative disease. Genes (Basel). 11, 1–24 (2020).

3. Slanzi, A., Iannoto, G., Rossi, B., Zenaro, E. & Constantin, G. In vitro Models of Neurodegenerative Diseases. Front. Cell Dev. Biol. 8, (2020).

4. Dawson, T. M., Golde, T. E. & Lagier-Tourenne, C. Animal models of neurodegenerative diseases. Nat. Neurosci. 21, 1370–1379 (2018).

5. Gitler, A. D., Dhillon, P. & Shorter, J. Neurodegenerative disease: Models, mechanisms, and a new hope. DMM Dis. Model. Mech. 10, 499–502 (2017).

6. Blanchard, J. W., Victor, M. B. & Tsai, L. H. Dissecting the complexities of Alzheimer disease with in vitro models of the human brain. Nat. Rev. Neurol. 18, 25–39 (2022).

7. Breslauer, D. N., Lee, P. J. & Lee, L. P. Microfluidics-based systems biology. Mol. Biosyst. 2, 97–112 (2006).

8. Ingber, D. E. Human organs-on-chips for disease modelling, drug development and personalized medicine. 23, 467–491 (2022).

9. Shuler, M. L. & Toh, C. A guide to the organ-on-a-chip. 2016, (2016).

10. Rio, J. A. & Ferrer, I. Potential of Microfluidics and Lab-on-Chip Platforms to Improve Understanding of “ prion-like “ Protein Assembly and Behavior. 8, 1–18 (2020).

11. Li, Y. et al. Microfluidics-based systems in diagnosis of Alzheimer’s disease and biomimetic modeling. Micromachines 11, 1–18 (2020).

12. Gagliano, O. et al. Synchronization between peripheral circadian clock and feeding-fasting cycles in microfluidic device sustains oscillatory pattern of transcriptome. Nat. Commun. 12, 1–12 (2021).

13. Luni, C. et al. High-efficiency cellular reprogramming with microfluidics. Nat. Methods 13, 446–452 (2016).

14. Giulitti, S. et al. Direct generation of human naive induced pluripotent stem cells from somatic cells in microfluidics. Nat. Cell Biol. 21, (2019).

15. Zambon, A., Zoso, A., Luni, C., Frommer, W. B. & Elvassore, N. Determination of glucose flux in live myoblasts by microfluidic nanosensing and mathematical modeling. Integr. Biol. (United Kingdom) 6, 277–288 (2014).

16. Michielin, F. et al. The Microfluidic Environment Reveals a Hidden Role of Self-Organizing Extracellular Matrix in Hepatic Commitment and Organoid Formation of hiPSCs. Cell Rep. 33, (2020).

17. Arias, E. R., Striebel, J., Busskamp, V. & Habibey, R. Microfluidics for Neuronal Cell and Circuit Engineering. doi:10.1021/acs.chemrev.2c00212.

18. Kothapalli, C. R. et al. A high-throughput microfluidic assay to study neurite response to growth factor gradients. Lab Chip 11, 497–507 (2011).

19. Sundararaghavan, H. G., Monteiro, G. A., Firestein, B. L. & Shreiber, D. I. Neurite growth in 3D collagen gels with gradients of mechanical properties. Biotechnol. Bioeng. 102, 632–643 (2009).

20. Peyrin, J. M. et al. Axon diodes for the reconstruction of oriented neuronal networks in microfluidic chambers. Lab Chip 11, 3663–3673 (2011).

21. Park, J. et al. Three-dimensional brain-on-a-chip with an interstitial level of flow and its application as an in vitro model of Alzheimer’s disease. Lab Chip 15, 141–150 (2015).

22. Taylor, A. M., Dieterich, D. C., Ito, H. T., Kim, S. A. & Schuman, E. M. Microfluidic Local Perfusion Chambers for the Visualization and Manipulation of Synapses. Neuron 66, 57–68 (2010).

23. Park, J. Y. et al. Differentiation of neural progenitor cells in a microfluidic chip-generated cytokine gradient. Stem Cells 27, 2646–2654 (2009).

24. Yan, X. 2018_Pnas_Si_Spe. Proc. Natl. Acad. Sci. 2017 (2017) doi:10.1073/pnas.

25. Gough, F. Human embryonic stem cell research in Ireland: Ethical and legal issues. Med. Law Int. 11, 262–283 (2011).

26. Takahashi, K. et al. Induction of Pluripotent Stem Cells from Adult Human Fibroblasts by Defined Factors. Cell 131, 861–872 (2007).

27. Zhang, Y. et al. Rapid single-step induction of functional neurons from human pluripotent stem cells. Neuron 78, 785–798 (2013).

28. Fantuzzo, J. A. et al. μNeurocircuitry: Establishing in vitro models of neurocircuits with human neurons. Technology 05, 87–97 (2017).

29. Middelkamp, H. H. T. et al. Cell type-specific changes in transcriptomic profiles of endothelial cells, iPSC-derived neurons and astrocytes cultured on microfluidic chips. Sci. Rep. 11, 1–12 (2021).

30. Tolomeo, A. M. et al. NGN2 mmRNA-Based Transcriptional Programming in Microfluidic Guides hiPSCs Toward Neural Fate With Multiple Identities. Front. Cell. Neurosci. 15, 1–13 (2021).

31. Kane, K. I. W. et al. Automated microfluidic cell culture of stem cell derived dopaminergic neurons. Sci. Rep. 9, 1–12 (2019).

32. Mcdonald, J. C., Duffy, D. C., Anderson, J. R. & Chiu, D. T. Review General Fabrication of microfluidic systems in poly (dimethylsiloxane). Electrophoresis 21, 27–40 (2000).

33. Miller, P. G. & Shuler, M. L. Design and demonstration of a pumpless 14 compartment microphysiological system. Biotechnol. Bioeng. 113, 2213–2227 (2016).

34. Kojic, S. P., Stojanovic, G. M. & Radonic, V. Novel cost-effective microfluidic chip based on hybrid fabrication and its comprehensive characterization. Sensors (Switzerland) 19, (2019).

35. Dimmeler, S., Ding, S., Rando, T. A. & Trounson, A. Translational strategies and challenges in regenerative medicine. Nat. Med. 20, 814–821 (2014).

36. Neužil, P., Giselbrecht, S., Länge, K., Huang, T. J. & Manz, A. Revisiting lab-on-a-chip technology for drug discovery. Nat. Rev. Drug Discov. 11, 620–632 (2012).

